# Stretch Induced Hyperexcitability of Mice Callosal Pathway

**DOI:** 10.1101/019190

**Authors:** Anthony Fan, Kevin Stebbings, Daniel Llano, Taher Saif

## Abstract

Memory and learning are thought to result from changes in synaptic strength. Previous studies on synaptic physiology in brain slices have traditionally been focused on biochemical processes. Here, we demonstrate with experiments on mouse brain slices that central nervous system plasticity is also sensitive to mechanical stretch. This is important, given the host of clinical conditions involving changes in mechanical tension on the brain, and the normal role that mechanical tension plays in brain development. A novel platform is developed to investigate neural responses to mechanical stretching. Flavoprotein autofluoresence (FA) imaging was employed for measuring neural activity. We observed that synaptic excitability substantially increases after a small (2.5%) stretch was held for 10 minutes and released. The increase is accumulative, i.e. multiple stretch cycles further increase the excitability. We also developed analytical tools to quantify the spatial spread and response strength. Results show that the spatial spread is less stable in slices undergoing the stretch-unstretch cycle. FA amplitude and activation rate decrease as excitability increases in stretch cases but not in electrically enhanced cases. These results collectively demonstrate that a small stretch in physiological range can modulate neural activities significantly, suggesting that mechanical events can be employed as a novel tool for the modulation of neural plasticity.

## 1. Introduction

Modulation of neuronal communications has traditionally been viewed as chemically induced (Kandel et al., 2012). However, recent evidence shows that mechanical cues influence many biological processes (Wang et al., 2009). Neurons are particularly sensitive to their mechanical micro-environment. For example, ion channels are mechano-sensitive, such that a high-enough stress induces structural changes in a protein complex, and hence a change in functionality (Sachs, 2010). In addition, growth of neurons can be induced by an applied stretch (Pfister et al., 2004). For example, dorsal root ganglion neurons are able to respond to a steady stretch and adjust their length to as far as a thousand times their original length. Application of stretch in the physiological range to frog neuromuscular junctions increases the frequency of spontaneous activities and the amplitude of evoked activities (Chen and Grinnell, 1995) within tens of milliseconds. Embryonic drosophila motor neurons actively maintain a rest tension of about a nano-Newton, and the loss of that tension prevents neurotransmitter vesicle clustering at the presynaptic terminal (Siechen et al., 2009). Finally, stretching the axon of the motor neurons by 5% for 30 minutes increases vesicle clustering by 200% (Siechen et al., 2009).

Central nervous system neurons are also subjected to large mechanical stretches and tension with varying rates during development, tumor growth and injuries, and brain swelling. It has been shown that tension is able to direct the growth of primary hippocampal neuron in 2-D culture (Lamoureux et al., 2002). Various groups have looked at the effect of traumatic strain, strain rate, and stress on cell death regulatory mechanism in brain slice preparations, brain slice culture, dissociated primary culture, and cell line culture (Morrison 3rd et al., 2011; Franze et al., 2013). However, the effects of mechanical stretches on long and short term synaptic functions remain unclear.

Although it has been speculated that mechanical forces play a role in brain morphology (van Essen, 1997) and function (Tyler, 2012), no direct observations have yet been made to demonstrate their effect on brain functionality. In this study, we show, using mouse brain coronal slice, that both a small stretch and a small stretch rate can substantially modulate evoked and spontaneous neural activities in the callosal pathway. We developed an experimental setup that allows us to apply a prescribed amount of stretch in one direction across the brain slice. Concurrently, the setup allows evaluation of spontaneous and electrically evoked neural activities. These activities are quantified by imaging flavoprotein autofluorescence (FA), which originates from the change of oxidation state of mitochondrial flavoproteins upon neuronal activation (Shibuki et al., 2003; Reinert et al., 2007).

## 2. Materials and Methods

### 2.1. Brain Slicing

One month old mice of both sexes were obtained from the in-house animal facility at Beckman Institute for Advanced Science and Technology at the University of Illinois. The animal was deeply anesthetized with ketamine (100 mg/kg) and xylazine (3 mg/kg) and a transcardial perfusion was subsequently performed with chilled oxygenated (95% O2/5% CO2) slicing solution [sucrose (234 mM), glucose (11 mM), NaHCO3(26 mM), KCl (2.5 mM), NaH2PO4·H2O (125 mM), MgCl2·6H2O (10 mM), and CaCl2·2H2O (0.5 mM)] at 4 °C. The mouse was then decapitated and its brain was removed from the skull. The aligned brain tissue was submerged in chilled slicing fluid in a vibration slicer. Obtained slices 600 μm thick were left in a warm (32 °C) artificial cerebrospinal fluid (aCSF) [NaHCO3 (26 mM), KCl (2.5 mM), glucose (10 mM), NaCl (126 mM), NaH2PO4·H2O (1.25 mM), MgCl2·6H2O (3 mM), and CaCl2**•**2H2O (1.1 mM)] bath for an hour before experiments. We have previously found that 600 μm thick slices prepared in a similar fashion are robustly viable (Llano et al., 2014). In this study, all data are obtained from the callosal pathway. All procedures were approved by the Institutional Animal Care and Use Committee at the University of Illinois. All animals were housed in animal care facilities approved by the American Association for Accreditation of Laboratory Animal Care. Every attempt was made to minimize the number of animals used and to reduce suffering at all stages of the study.

### 2.2. Stretching Mechanism

Slices were made stationary by a micro-vice on each side of the perfusion stage as depicted in Fig. 1A-C. Micro-channels enable aCSF [NaHCO3 (26 mM), KCl (2.5 mM), glucose (10 mM), NaCl (126 mM), NaH2PO4•H2O (1.25 mM), MgCl2**•**6H2O (2 mM), and CaCl2·2H2O (2 mM)] to perfuse the slice at a constant flow rate with no apparent waste accumulation in the recording chamber. Each micro-vice is connected to the frame of the recording chamber. A power screw mechanism controlled by a stepper motor is used to actuate the micro-vices towards or away from each other. Each clockwise turn of the screw would lead to a linear retraction of 1/80 inches and vice versa. Geometries were optimized so that the whole device could be fit under the fluorescence microscope used for imaging during the experiment.

**Figure 1:**
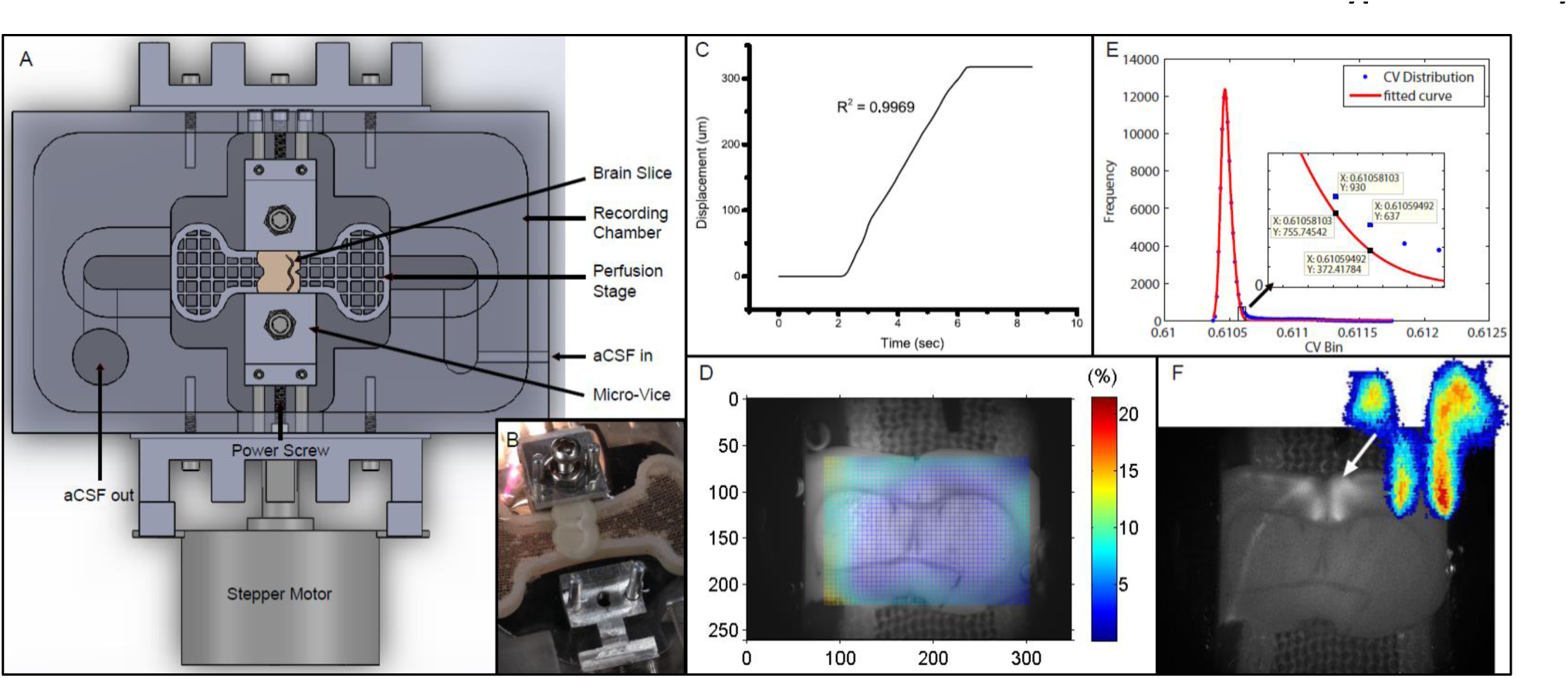
Experimental setup and data analysis methods. (A) Schematics depicting the entire setup. Micro-vices are actuated by a power screw mechanism in which a rotation of the screw will result in a uniaxial translation. (B) Picture of device with brain slice secured on one end. (C) Displacement vs. time plot depicting the stretch applied during the experiment. R^2^ value describes a linear fit to the increasing region. The displacement is 317.5 μm over 4 seconds, giving a strain rate of 79.4 μm/s. (D) Digital image correlation (see methods) was used to obtain displacement fields from videos. The color map here plots the effective strain (Eq. 2) over the brain slice preparation. (E) I_cv_ of all pixels are plotted as a histogram in blue circles. Red line depicts a 2-term Gaussian fit. The right hand side is magnified in the subplot. Threshold value based on Eq. 3 is sandwiched between the 2 data labels. (F) Background image was obtained by performing a standard deviation projection. The heat map was generated by plotting 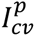. Sparse outliers away from the main area were also trimmed.

### 2.3. Flavoprotein Autofluorescence Imaging

FA was used in this study to track neuronal activities. FA signal is intrinsic and its consistency is confirmed to be reliable for several hours (Shibuki et al., 2003). The method provides a rapid assessment of neuronal activation and carries several advantages compared to other imaging techniques that involve dye loading. Dye loading (e.g., calcium-sensitive or voltage-sensitive dyes) creates substantial changes in signal strength over time and potential heterogeneities over the tissue, depending on dye uptake. Electrophysiological measurements are of limited use in the current experiments because upon the induction of stretch, slippage might occur between the electrode-tissue interface. In this study, the slice was put under an epifluorescence microscope [Olympus BX51 with a Prior Lumen 200 light source] using anUMNIB Olympus filter cube (470- to 490-nmexcitation, 505-nm dichroic, and 515-nm emission long pass) and other supplementary optics (Theyel et al., 2011). Images were acquired with a CCD camera [QImaging EXi with Firewire interface] at 4 frames per second. Image analyses were done in ImageJ and MATLAB.

### 2.4. Electrical and Chemical Stimulation

In all experiments, 800 nM of SR95531 (Tocris Bioscience, Bristol, UK), antagonist of the neuroinhibitor GABAA, was added to the aCSF. SR95531 lowers the activation barrier for neuronal firing to facilitate experimental readouts. To stimulate, we used in-bath tungsten electrodes that did not directly contact the slice. One-second-long electrical pulse train stimulations were applied every 10 seconds for 50 repetitions in all experiments. Each pulse train has a step profile of 1-6 mA for 2 ms at 40 Hz.

### 2.5. Strain field calculation

Digital image correlation (DIC) analysis (Jones et al., 2014) was performed on the acquired images to find the corresponding displacement vector at each subregion. The displacement values were then formulated into a 2-D strain tensor field (Fig. 1D), given the plane stress assumption, using the following strain-displacement relationship:

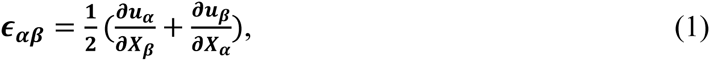

where *α*, *β* = 1, 2 define the in-plane coordinates. Effective strain (∊_e_), reported as strain in text, can be defined using the Einstein notation as:

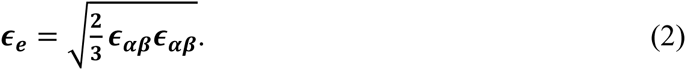

### 2.6. Quantification of Evoked Activation

We used the coefficient of variation, *I*_*cv*_, (standard deviation over mean) of temporal intensities of each pixel to find the active pixels in our image sequences. To determine the threshold for quantification of activation area, we plot the *I*_*cv*_ distribution into a histogram, *H*[*I*_*cv*_], and fit the profile to a two-term Gaussian function, *G*(*I*_*cv*_). The right tail (with larger values) is expected to deviate from the normal distribution, and we use the point of deviation as our *I*_*cv*_ threshold value (Fig. 1E). The point of deviation, 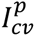 is defined as:

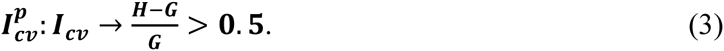

To quantify the time profile of activation of the activated region (Fig. 1F), spatial averaging of intensity is used. The change in activation is quantified by 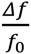, defined as the change in fluorescence over baseline fluorescence.

For spontaneous activities quantification during stretching, the adjusted intensity is defined as:

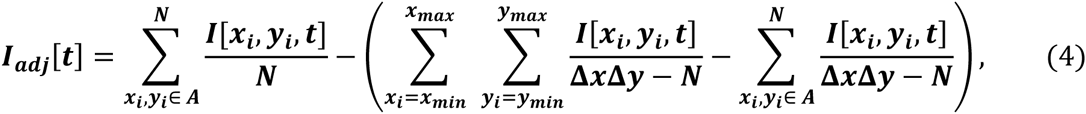

where A denotes the activation area. *x*_*max*_, *x*_*min*_, *y*_*max*_, and *y*_*min*_ are the limit points of the smallest rectangle enclosing the entire activation area. Δ*x* = *x*_*max*_– *x*_*min*_. Δ*y* = *y*_*max*_ − *y*_*min*_. The first term in Eq. 4 denotes the signal. The second term denotes background intensity.

## 3. Results

### 3.1. Flavoprotein Autofluorescence before Stretch and after Stretch

Using the platform we developed (Fig. 1A), the brain slice was gently gripped and tensed until no edge slippage was observed (see Fig. 1B and Materials and Methods for detailed description). As baseline measurement, we took a fluorescence video of 500 seconds long recording the response of the slice to 50 stimulation pulse trains, each 10 seconds apart. The slice was subsequently stretched by 317.5 μm (or 4.2% of total the length of the slice) over 4 seconds (Fig. 1C). Using the DIC methods (see Materials and Methods), effective strain at responsive area was found to be 2.5% (Fig. 1D), while edge-to-edge global effective strain is 4.2 %. The stretch was maintained for 10 minutes and then entirely released. *No measurement was made while the slice was being held.* After the slice was released from stretch, we performed the baseline measurement again. This stretch-baseline-cycle was repeated for 4 more times. This paradigm was applied to 4 independent slices from 4 animals.

To ensure that the extra excitability is not due to sensitization to electrical stimulations or degeneration of tissue health, control slices were subjected to the same series of manipulation, without the stretch. The control paradigm was applied to 3 independent slices from 3 animals.

### 3.2. Stretch Effects on Excitability

Excitability here is defined as the ratio of number of times the slice responds (detected using FA) to the number it is electrically stimulated. Thus excitability gives the probability of FA response. We found that in all slices undergoing the stretch-baseline-cycle (n=4), excitability increases after every cycle to ultimately 3 times of the original probability (Fig. 2A). It thus seems that the slice “remembers” its past history of stretch, and its current excitability results from a cumulative effect of its past stretches. Normalized baseline measurements from the control group (n=3) are plotted next to stretched group. We fit a linear regression to each experiment individually, and the averaged slope is reported in Fig. 2B comparing the slope in stretched and control slices. We found that the stretched slices showed a significant increase in excitability compared to control slices (p=0.013).

**Figure 2:**
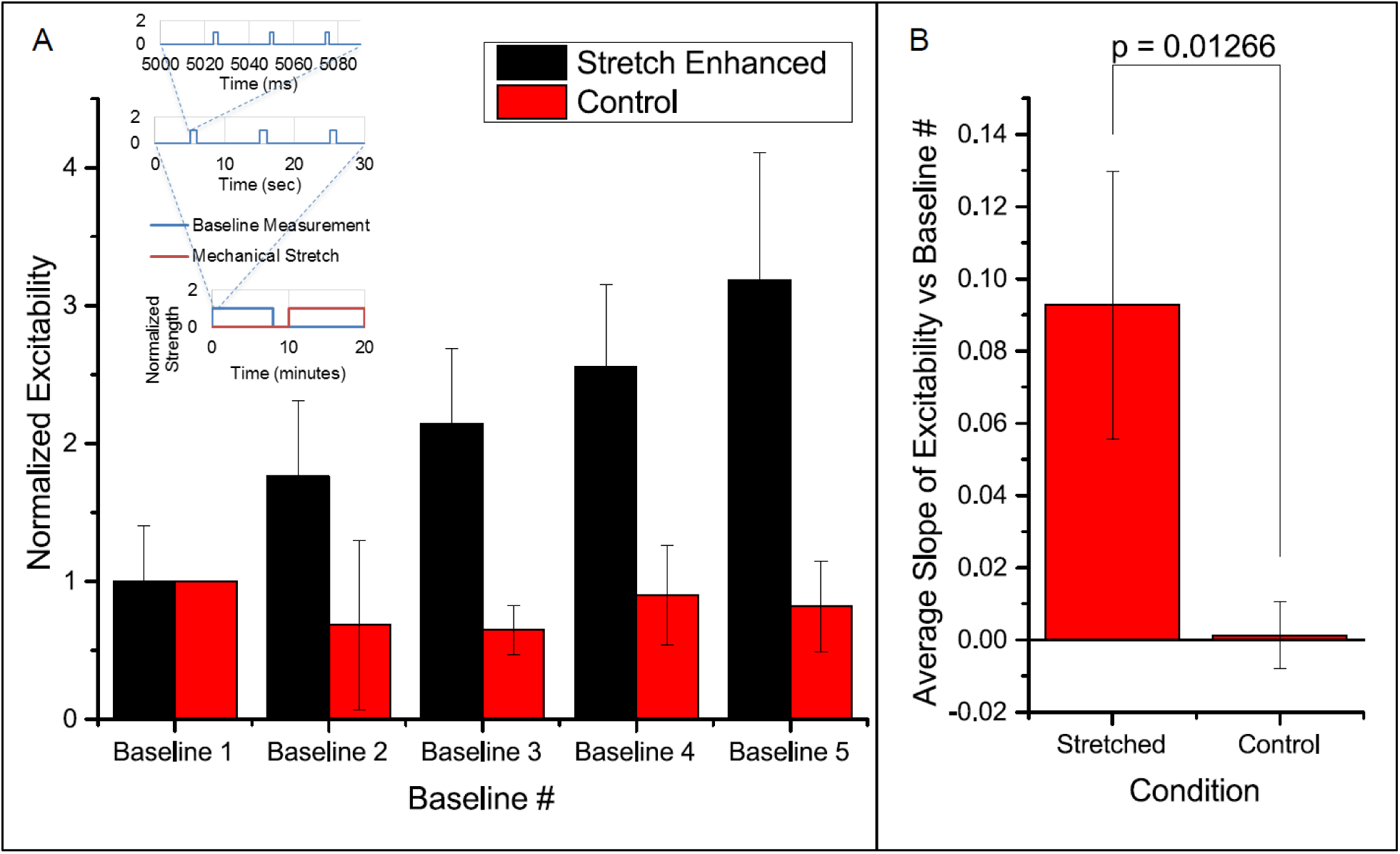
Enhanced excitability of callosal pathway by stretch alone and the 2 control schemes. (A) Normalized probability of response of stretched group and control group. The stretched data set here is individually normalized to the average probability of the first baseline. This serves to show the variation in the starting probability of response, as given in the error bar. All subsequent column plots are normalized to the first baseline probability within individual experimental data set, leading to an error bar magnitude of 0. Subfigure includes a schematic of the paradigm for one cycle. (B) The slopes of excitability vs baseline # in each independent data set are averaged and reported here. This serves to compare the increase in excitability of the stretched and control groups. All error bars in SD. P-value obtained from 2-tail t-test with unequal variance.

### 3.3. Stretch Effects on Spatial and Temporal Activation

We note that the area of activation of the slice fluctuates in the stretched group (Fig. 3A). Further looking into the spatial distribution and the strength of variation in the activation area, we observed a shift in the location of maximum activity in the stretched group. An example is shown in Fig. 3A, with the stretched group at the top row and the control group at the bottom. It is thus possible that the stretch applied and released in between the 2 baseline measurements can lead to a spatial reconfiguration of excitability.

**Figure 3:**
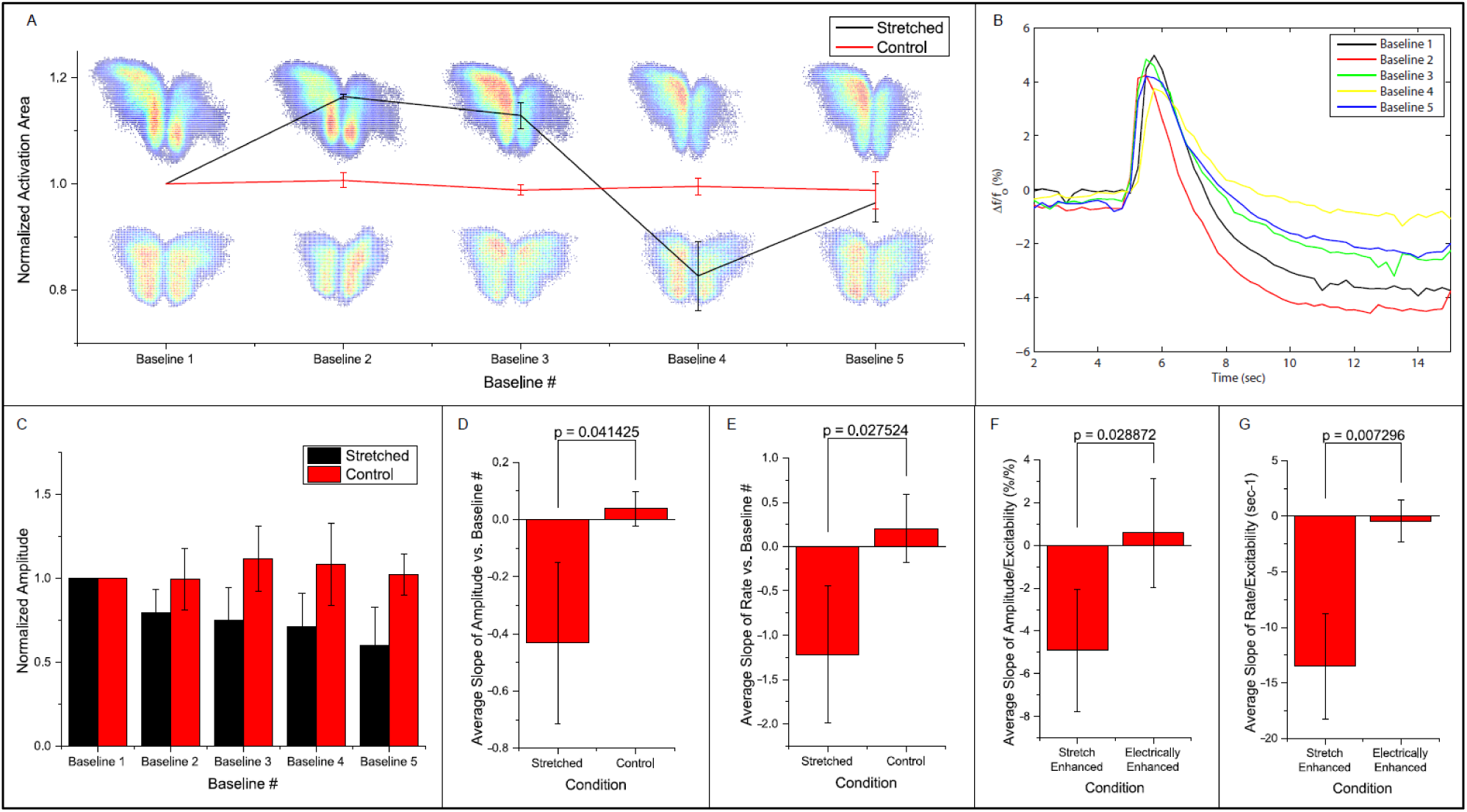
Fluorescence intensities are averaged over the calculated activation areas. (A) Spatial activities in subsequent baseline measurements. Normalized activation area of the stretched and control groups are plotted. Two examples of activation areas: stretched on top, and control at the bottom. (B) An example of the 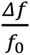 signal in one experiment. Each line denotes the first peak in respective baseline measurement. (C) Normalized average amplitudes are plotted for the stretched and control groups. Slope of (D, F) amplitude and (E, G) rate vs baseline measurement number and excitability from each experiment is averaged and compared. All error bars in SD. P-values obtained from 2-tail t-test with unequal variance.

Temporal activation profile, 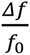, is plotted against time. The amplitude of each fluorescence peak is obtained by subtracting the value preceding the stimulation to the peak 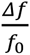 value. The first activation amplitude, i.e. the first electrical stimulation after the stretch is released, in each baseline measurement is found to be steady (Fig. 3B). However, the average activation amplitude is found to decrease after each stretch-baseline cycle. The control amplitudes remain unchanged (Fig. 3C). We fit a linear regression to the amplitudes in each experiment individually, and the averaged slope is reported in Fig. 3D. All stretched slices show significant negative slope compared to controls (p = 0.041) and a significant negative slope of the temporal activation rate (p = 0.028, Fig. 3E).

The attenuations in the signal appear to correlate with the excitability. This leads to the question whether the attenuation of activation amplitude and rate is simply due to the increase in excitability or is actually induced by the stretching paradigm. To address this question, we artificially increased the probability of FA response by increasing the applied electrical stimulation (n=4). A linear regression is fit to the amplitude/rate vs. excitability profile in individual experiment separately (Fig 3F & G). The average slope shows that the activation amplitude and rate do not change with electrical stimulation alone. Thus, we conclude that the attenuation of the signals is due to the stretching paradigm rather than a consequence of electrical stimulation, suggesting that the stretching manipulation might have lowered the energy barrier for synaptic activation

### 3.4. Spontaneous Activities due to Stretch Alone

A separate slice was stretched with time (Fig. 4A) in a stepwise fashion in 3 cycles with 2 μM of SR95531. There was a 5 minute time gap between each cycle. No electrical stimulation was applied here. The slice was imaged during the entire span of the cycles and FA data were analyzed as before. The 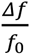 plots show a close correspondence between stretching impulse and induced flavoprotein activities in all three cycles of stretch (Fig. 4A). No activity was observed during the relaxation steps. Out of the 11 stretching steps, 6 (50%) led to an immediate FA response; two others led to FA responses within 13 seconds (Fig. 4B). The results suggest that a brain slice is sensitive to both maintained stretches and stretch impulses.

**Figure 4:**
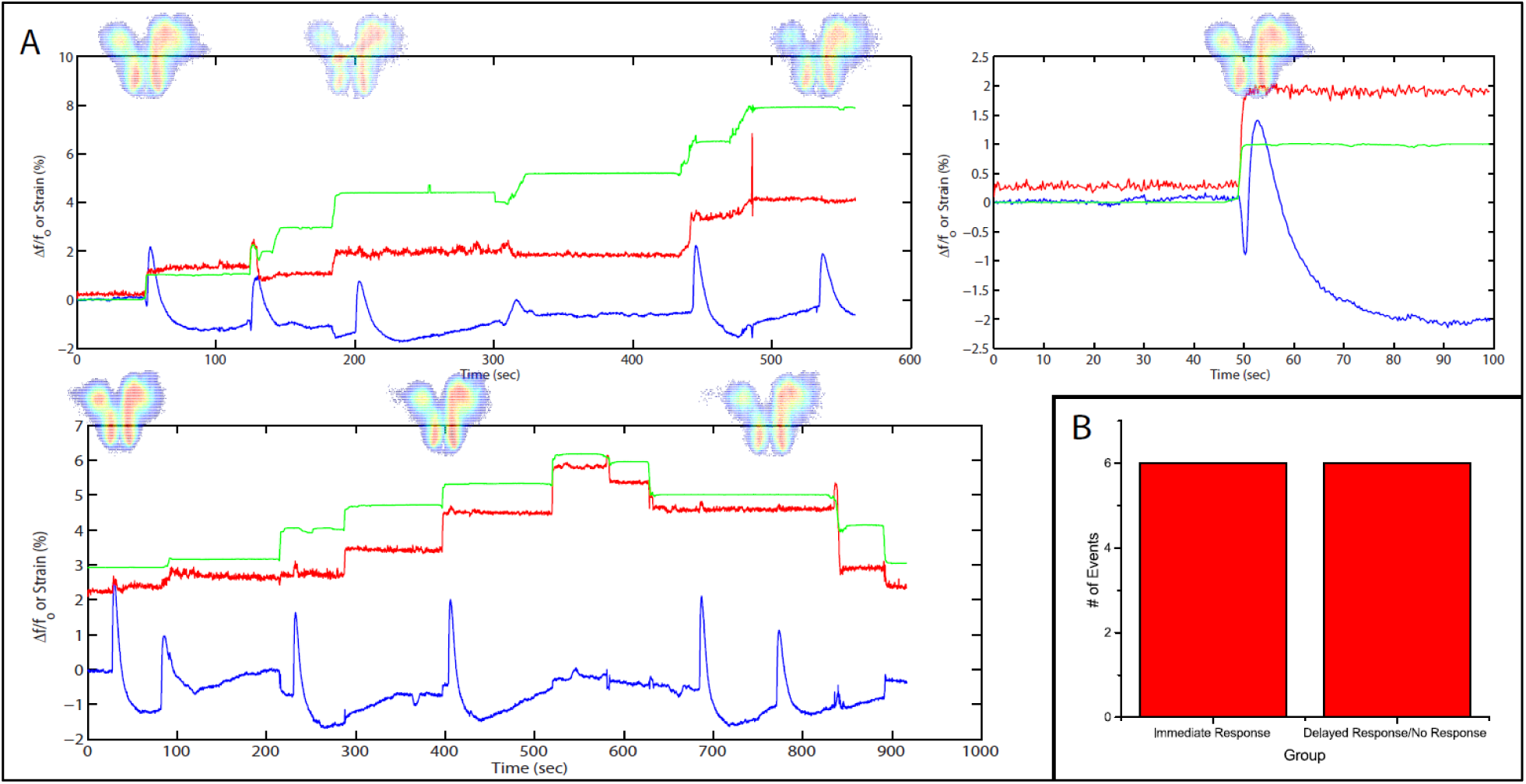
Stretch Effects Alone without Electrical Stimulation. (A) Processed average intensities are plotted against time. Red lines denote effective strain level at activation area. Green lines denote effective strain level globally, defined by the position of the micro-vices. Activation area examples are placed aligned to respective peaks. (B) All fluorescence occurrences are grouped into 2 groups: 1) immediate firing after stretch application, 2) delayed firing or no firing before the next stretch.

## 4. Discussion

The probability of neural response to electrical stimulation can be increased by applying a stronger electrical stimulus or by applying the stimulus more frequently (Bikson et al., 2004). Long-term potentiation (LTP) has been shown to be induced by a tetanic stimulation in which frequency is usually in the range of 100Hz (Malenka and Bear, 2004). Long-term depression (LTD), on the other hand, can be induced by a low frequency stimulation in the range of 10Hz (Malenka and Bear, 2004). The actual frequency that is required to induce plasticity is pathway-dependent and species-dependent (Malenka and Bear, 2004). At intermediate frequencies, synaptic transmission is usually memoryless, i.e., the same electrical stimulus will give the same probability/strength of neural responses regardless of what the pathway was subjected to before (Bliss and Lomo, 1973).

Here we show that by maintaining a small stretch for 10 minutes, we can increase this probability of neural response. Note that this is not the same as LTP, since LTP describes the changes in strength of postsynaptic response, whereas we are reporting excitability. Nonetheless, in both LTP and the stretch-enhanced excitability, the electrically evoked activity has a higher probability in surpassing the activation threshold.

It is not yet known whether the changes in probability, amplitude or rate of the FA response correlate with similar change in the underlying electrophysiological response. However, it has been shown that there is a strong correspondence in the strength of the 2 signals under normal conditions (Shibuki et al., 2003; Reinert et al., 2004; Llano et al., 2009). It has also been shown that only 15% of the FA signal is left when postsynaptic glutamate receptors are blocked (Llano et al., 2012; Reinert et al., 2004). Therefore, it is likely the increase in excitability we see here are due to direct/indirect enhancement in postsynaptic activity—either as a result of more vesicles being released from the presynaptic terminal upon the same stimulation, or the mechanism of receiving the vesicles has become more efficient.

Previous finding reports that a 5% stretch in embryonic Drosophila axons of motor neurons results in a 200% increase in neurotransmitter vesicle clustering at the presynaptic terminal of neuromuscular junctions. It takes about 20 minutes for the increased clustering to occur (Siechen et al. 2009; Ahmed et al. 2012). In our case, the stretch was held for 10 minutes. Arguably, the difference could be attributed to the different species and the different neural structures (neuromuscular vs. neural-neural). It is thus possible that the increased clustering of synaptic vesicles can lead to a higher docking ratio, more frequent spontaneous release, and larger release upon stimulation (Dobrunz, 2002). This could explain the higher excitability upon completion of a stretch-baseline cycle.

The increased response due to stretch could be because of postsynaptic modifications, in particular, that stretch results in increased clustering of neurotransmitter receptors (AMPA and/or NMDA) in the postsynaptic density, similar to LTP (Bear et al., 1987). This increases the probability of neuronal activity due to a given electrical stimulation and hence a given presynaptic release of neurotransmitter. It has been shown that integrin activities (to be discussed) lead to phosphorylation of CamKII (Charrier et al., 2010), known to be critical in LTP (Lisman et al., 2012).

Cell-cell adhesion protein molecules facilitate force transfer in neighboring cells by acting as an anchor (Wang et al., 2009). Many of these protein complexes are found in neural pathways (Mobley et al., 2009). Increased mechanical tension can lead to a structural change in these molecules leading to downstream signaling cascades. One study has attributed the high stretch sensitivity of motor nerve terminals to integrin, a cell-cell adhesion protein (Chen and Grinnell, 1995). In that study, both the evoked amplitude and the frequency of spontaneous activity increase significantly with the application of stretch on frog muscle. The increase is linear with respect to the magnitude of the stretch applied. The known integrin inhibitors, such as peptides containing the Arg-Gly-Asp (RGD) sequence, suppressed the stretch sensitivity. Enhancement of transmitter release occurs within a few milliseconds of the stretch application, which might affect the stretch-rate sensitivity of neuronal response, i.e. increased responsiveness at faster rate of stretch.

Stretch or tension can also increase ionic conductances, similar to peripheral mechanotransducers (Sachs, 1991), either pre- or post-synaptically. Collectively, or individually, alterations in ionic conductances could lower the activation barrier for action potentials or increase post-synaptic sensitivity to presynaptic activity. The change of conductance due to stretch is expected to be instantaneous. Thus this mechanism should also result in strain-rate sensitivity, possibly providing an explanation for the similar sensitivity we observed.

There are also clinical implications to this work. Stretching of axons is seen in a large number of clinical conditions, including hydrocephalus, traumatic brain injury and other forms of brain injury leading to edema (e.g., stroke, tumors, infection). The current results imply that even small amounts of stretch can lead to major changes in neuronal function, and may explain hyperexcitability phenomena, such as seizures, that can be seen during these states. Future work will clarify the mechanisms of the changes we observed, and may therefore lead to novel therapeutics to deal with these stretch-induced hyperexcitability.

## Acknowledgments

This work was funded by the National Science Foundation (NSF) Grant 0965918 IGERT at UIUC: Training the Next Generation of Researchers in Cellular and Molecular Mechanics and Bio-Nanotechnology.

## Contributions

AF, KS, DL, TS designed research. AF, KS performed research. AF contributed unpublished reagents/analytic tools. AF, TS analyzed data. AF, TS wrote the paper. All authors read and reviewed the paper.

